# Calreticulin modulates the infection process and nodule organogenesis in the *Phaseolus vulgaris-Rhizobium* symbiosis

**DOI:** 10.64898/2026.04.09.717402

**Authors:** Yolanda Ortega-Ortega, Janet Carrasco-Castilla, Marco A. Juárez-Verdayes, Noreide Nava, Jorge Solis-Miranda, Ronal Pacheco, Carmen Quinto

**Affiliations:** Departamento de Biociencias y Agrotecnología, Centro de Investigación en Química Aplicada-SECIHTI, Saltillo, Coahuila, México; Centro de Estudios Científicos y Tecnológicos No. 17, Instituto Politécnico Nacional, León, Guanajuato, México; Departamento de Ciencias Básica, Universidad Autónoma Agraria Antonio Narro, Calzada Antonio Narro, Saltillo, Coahuila, Mexico; Departamento de Biología Molecular de Plantas, Instituto de Biotecnología, Universidad Nacional Autónoma de México, Cuernavaca, Morelos, México; Instituto de Bioquímica Vegetal y Fotosíntesis, Consejo Superior de Investigaciones Científicas (CSIC)-Universidad de Sevilla, Sevilla, España; Programa de Ecología Genómica, Centro de Ciencias Genómicas, Universidad Nacional Autónoma de México, Av. Universidad s/n, Chamilpa 62210, Cuernavaca, Morelos, Mexico

**Keywords:** Calreticulin, nodulation, *Phaseolus vulgaris*, nitrogen-fixation

## Abstract

Calreticulins are multifunctional proteins involved in calcium homeostasis, protein folding, and cellular signaling. In common bean (*Phaseolus vulgaris*), the molecular mechanisms that regulate infection and nodule development remain incompletely understood. The main objective of this study was to characterize the role of the calreticulin gene *PvCRT08* during infection and nodulation processes. We first analyzed the calreticulin gene family in the *P. vulgaris* genome and identified three members, with *PvCRT08* showing the highest transcript accumulation in roots and after inoculation with rhizobia. Spatial and temporal promoter analyses in transgenic composite bean roots revealed *PvCRT08* activity in root hairs and in infected cells and vascular bundles of mature nodules. RNA interference (RNAi)-mediated *PvCRT08* down-regulation in transgenic roots increased the number of infection threads and enhanced nitrogen fixation efficiency, leading to the formation of larger and more functional nodules, although total nodule number was unaffected. In contrast, overexpression of *PvCRT08* impaired infection thread progression, reduced the expression of key nodulation marker genes (*PvCyclin* and *PvNIN)*, decreased nodule number, and diminished nitrogen fixation capacity. These findings identify *PvCRT08* as a key regulatory component of early infection events and nodule development in common bean. Furthermore, the study provides new insights into the molecular control of symbiotic efficiency and highlights *PvCRT08* expression is critical to optimize the equilibrium between infection efficiency and nodule functionality.

## Introduction

Nitrogen is essential for all living organisms as it is a major component of amino acids and nucleic acids. Although the Earth’s atmosphere is composed of 78.1% diatomic nitrogen gas (N_2_), plants are unable to use this form of nitrogen, so most farmers use chemical nitrogen fertilizers to improve crop productivity. However, the excessive use of nitrogen fertilizers causes serious environmental damage **[1,2]**. The nitrogenase enzyme and the ability to fix N are exclusively prokaryotic. There are free-living and symbiotic nitrogen-fixing bacteria, the latter associated with the root of legumes, some of which belong to the Rhizobiacea family **[3,4].** Common bean (*Phaseolus vulgaris* L.) is an important legume crop as it has been part of the human diet since ancient times. It belongs to the family Fabaceae, subfamily Papilionoideae, tribe Phaseoleae, subtribe Phaseolinae, genus *Phaseolus* and *Phaseolus vulgaris* species **[5].**

Nodule formation is the result of the symbiotic interaction between legumes and nitrogen-fixing soil bacteria of the family Rhizobiaceae (called rhizobia). This process begins through a highly specific molecular dialog between both symbionts. The roots of the host plant exude flavonoid compounds into the rhizosphere, which attract bacteria to the apical region of the root hairs and activate the bacterial gene transcription factor NodD. This factor controls the expression of nodulation genes in rhizobia. These nodulation genes are involved in the synthesis and secretion of specific lipo-chito-oligosaccharides, also known as Nod Factors (NFs) **[6,7].** The NFs are recognized by receptors in the root hair, triggering morphological and physiological changes such as root hair curling, which traps bacteria, and initiates the formation of the infection thread (IT). The IT is a tubular invagination of the host cell wall and plasma membrane, the lumen of which is filled with bacteria. The IT grows within the root hair toward the cortex cells and eventually branches within the nodule primordium cells from which the nodule develops, ultimately leading to biological nitrogen fixation. Both infection thread formation and nodule organogenesis processes are regulated by various biochemical signals, mainly calcium spiking and the decoding of this signal **[8–11].**

Calreticulin (CRT) is a protein that is ubiquitously expressed in all multicellular eukaryotes, such as humans, nematodes and plants. Several studies indicate that CRT is found primarily in the endoplasmic reticulum (ER), but it can also be found in the nuclear envelope and nucleus. The protein participates in several intra and extracellular processes, such as the regulation of intracellular Ca^2+^ homeostasis, apoptosis, the control of cell adhesion and it exhibits chaperone activity in the folding of newly synthetized proteins in the ER **[12–14].**

The CRT is a chaperone protein with a molecular weight of 46 kDa, highly conserved and located mainly in the ER. The protein contains three distinct structural and functional domains: a globular N-domain, a proline-rich P-domain, and a polyacid C-domain. The N domain includes two well-conserved motifs (KHEQKLDCGGGYVKLL and IMFGPDICG) and a signal peptide for the ER. The middle third, or P-domain of the CRT, is a proline-rich segment responsible for the high affinity (Kd=1M), and low capacity (1 mol Ca^2+^/ 2mol of protein) of the Ca^2+-^binding site. It contains two sets of three sequence repeats (A and B) implicated in the chaperone function. These sequences are highly conserved among the CRTs of various animal and plant species. In plants, the A repeat has the structure of PXXIXDPXXKKPEXWDD while the consensus sequence for the B repeat is GXWX-AXXIXNPXYK; in contrast, the repeated sequences in animals are PXXIXDPDAXKPEDWDE and GXWXXPXIXNPXYK, respectively. The C-domain contains many negatively charged residues, is highly acidic, and binds Ca^2+^ with a relatively high capacity but low affinity (25 mol of Ca^2+^/mol of protein). The C-domain ends with the ER recovery signal KDEL (Lys-Asp-Glu-Leu) in animals, and the sequence HDEL (His-Asp-Glu-Leu) in plants **[12,13,15].**

Earlier, using a phosphoproteomic approach, we analyzed the abundance of a phosphorylated calreticulin (phytozome accession number Phvul.007G200800, PvCRT08) in *P. vulgaris* roots in response to inoculation with *Rhizobium etli*. The results obtained showed that these levels increased significantly up to 24 h (Ortega-Ortega, unpublished data). Based on these results, we decided to explore the role of this protein in the bean-rhizobia symbiosis through reverse genetics. To achieve this, we first analyze whether the gene of interest is part of a gene family and then determined the levels of accumulation of the corresponding transcript, in response to inoculation with rhizobia. We also examined the promoter activity of the gene at different stages of nodulation. Regarding the functional analysis of the gene, knocking-down and overexpression approaches were performed to resolve this issue. The results obtained indicate that the promoter activity of *PvCRT08* was detected in the root hairs inoculated with rhizobia and in the vascular bundles of mature nodules. The participation of *PvCRT08* in the common bean symbiosis with *Rhizobium tropici,* using these approaches revealed that this protein plays a regulatory role in both infection and nodulation processes.

## Materials and Methods

### Plant growth conditions and Rhizobia inoculation

Seeds of *P. vulgaris* cv. Negro Jamapa (obtained from the local farmers’ markets, Cuernavaca, Mexico) were surface-sterilized and germinated at 28°C for 2 days in the dark. At 2 days post-germination, the apex of root was sectioned from seedlings and frozen in liquid nitrogen. Separately, the root hairs were gently broken off from roots without apex using a magnetic stir bar in a stainless-steel tank with liquid nitrogen. The bared roots were separated by filtering through a sieve, and the root hairs were collected in liquid nitrogen. All the material was stored at −70 °C until subsequent RNA extraction and reverse-transcription quantitative PCR (RT-qPCR) assays.

Composite common bean plants were generated according to the protocol developed by Estrada-Navarrete et al **[16].** Hairy roots (10–13 days post-emergence) were generated using *Rhizobium rhizogenes* strain K599 **[17].** Transgenic composite plants carrying the corresponding construct were observed under epifluorescence microscopy to confirm the presence of the reporter gene (GFP or TdTomato), and untransformed roots were removed. Composite common bean plants were planted in pots with vermiculite and inoculated with 20 mL of *R. tropici*–GUS or *R. tropici*–DsRed suspension at an optical density at 600 nm (OD600) of 0.05 (undiluted suspension had OD600 = 0.8–1.0). Plants were grown under greenhouse conditions with a controlled environment (26–28 ◦C, 16 h light:8 h dark) and were watered with B&D medium **[18]**.

### Identification of *PvCRT* sequences, and relative transcript accumulation in different organs and tissues

Using the Phvul.007G200800 as template, searches for CRTs in the Phytozome v12 database (http://phytozome.jgi.doe.gov/pz/portal.html) **[19]** were performed for *P. vulgaris, Glycine max, Medicago truncatula and Arabidopsis thaliana. Oryza sativa, Marcanthia polymorpha, Selaginella moellendorffii* and *Homo sapiens* protein sequences homologous to CRT were identified and downloaded from the PlanGDB (http://www.plantgdb.org/), MarpolBase (https://marchantia.info/nomenclature/) and NCBI (https://www.ncbi.nlm.nih.gov/) databases, respectively (**S1 Table**). Multiple sequence alignment of the CRT amino acid sequences was performed using MUSCLE Geneious prime software. A phylogenetic tree was generated by the maximum likelihood method based on the JTT matrix and FreeRate with 3 categories using Geneious prime software version 2020.2.4 from 10,000 bootstrap replicates. The *P. vulgaris* Genome Expression Atlas (PvGEA, http://plantgrn.noble.org/PvGEA/) was used in order to estimate the *PvCRT* transcript accumulation profile from different organs and tissues of *P. vulgaris* analyzed by RNA sequencing and expressed as reads per kilobase of transcript per million mapped reads.

### RT-qPCR assays

High quality RNA was isolated from frozen tissues using TRIzol®Reagent (Invitrogen, Life Technologies, USA) following the manufacturer’s recommendations. RNA integrity and concentrations were determined by electrophoresis and NanoDrop (2000c, Thermo Scientific) spectrophotometry, respectively. Genomic DNA contamination from RNA samples was removed by incubating the samples with RNase-free DNase (1 U·µL^−1^) at 37 °C for 15 min and then at 65 °C for 10 min. RT-qPCR assays were performed using an iScript™ One-step RT-PCR Kit with SYBR®Green (Bio-Rad) from DNA-free RNA samples, according the manufacturer’s recommendations, in a LightCycler®Nano cycler (Roche). RNA template concentration was 40 ng (10 ng·µL−1) in each reaction. DNA-free RNA samples were used as a control to confirm the absence of DNA contamination. Relative expression values were calculated using the formula 2−ΔCT, where cycle threshold value (ΔCt) is the cycle threshold (Ct) of the gene of interest minus the Ct of the reference gene **[20,21]**. RT-qPCR data were generated from at least two biological replicates with three independent plants each one, together with three technical replicates. The gene encoding elongation factor 1-alpha (EF1α) was used as a reference gene to normalize the experimental data. The genes analyzed are listed in **S2 Table**, together with their specific oligonucleotides used.

### Plasmid construction

To develop a p*PvCRT08*:GUS-GFP construct, a 2086-bp fragment upstream of the *PvCRT08* translation start site was amplified using bean genomic DNA and primers p*PvCRT08*-Up and p*PvCRT08*-Lw (**Table S6**) and cloned into vector pENTR/SD/D-TOPO (Invitrogen). The Gateway LR reaction was performed between the entry vector pENTR/SD/D-TOPO-*PvCRT08* and the destination vector pBGWFS7.0 **[22]** according to the manufacturers’ instructions (Invitrogen). Control transgenic roots harbored a cassette with no promoter sequences upstream of the GFP-GUS sequence.

To generate the RNAi construct, a fragment corresponding to the 3’-untranslated region of the *PvCRT08* gene was amplified from cDNA isolated from common bean roots at 2 days post germination using the following primers: *Pv*CRT08-RNAi-Up (5’-TAGGCAGTTTGCAAAACGGAT-3’) and *Pv*CTR08-RNAi-Lw (5’-AGATTGAAACAAAAGCTCTCCAGC-3’). The resulting PCR product was cloned into pENTR/D-TOPO (Invitrogen) vector and transformed by heat-shock into *Escherichia coli* TOP10 chemically competent cells. The recombination into the destination vector pTdT-DC-RNAi **[23]** was performed with the LR clonase, using the Gateway System (Invitrogen). The appropriated orientation of the insert was confirmed by PCR and sequencing using the *PvCRT08*-RNAi-Up primer for the pTdT-*PvCRT08*-RNAi plasmid together with WRKY-5-Rev (5’-GCAGAGGAGGAGAAGCTTCTAG-3’) or WRKY-3-Fwd (5’-CTTCTCCAACCACAGGAATTCATC-3’) primer. As a control, a truncated and irrelevant sequence from *A. thaliana* pre-mir159 (kindly provided by Dr. José Luis Reyes), lacking the target sequence of miR159 (ACAGTTTGCTTATGTCGGATCCATAATATATTTGACAAGATACTTTGTTTTTCGATAGATCTTGATCTGACGATGGAAGTAGAGCTCTACATCCCGGGTCA), was cloned into the pTdT-DC-RNAi vector. The correct orientation of the sequence in the construct was confirmed by DNA sequencing.

To construct an overexpression vector for *PvCRT08*, the 1263-bp ORF of *PvCRT08* (Phvul.007G200800) including the 5’ -untranslated region (56 bp) and a 3’-untranslated (124 bp) fragment was isolated from *P. vulgaris* cDNA. This region was amplified from *P. vulgaris* root cDNA at 2 days post-germination using *PvCRT08*-OE-Up and *PvCRT08*-OE-Lw primers (**Table S2**). The fragment was cloned into the pENTR/D-TOPO vector (Invitrogen) and sequenced. The resulting pENTR-*PvCRT08* plasmid was recombined into the binary vector pH7WG2tdT under the control of the constitutive 35S promoter **[24]**. Briefly, this vector was derived from vector pH7WG2D; the cassette pEgfpER and the 35S terminator were respectively replaced by a TdTomato reporter (red fluorescent protein) obtained from the pTd-DC-RNAi vector and an E9 terminator. Empty pH7WG2tdT vector was used as the control.

### Promoter activity analysis

Composite *P. vulgaris* plants harboring p*PvCRT08*:GUS-GFP were transferred into pots of vermiculite and each plant was inoculated with 20 mL of *R. tropici*–DsRed suspension (with an OD_600_ of 0.05). Roots and nodules were collected at 3, 5, 7 and 21 dpi (days post-inoculation). Samples were histochemically analyzed for GUS activity according to the method of Jefferson **[25]** and images were acquired with a Retiga 4000R CCD camera coupled to a Nikon TE300 inverted microscope. Promoter activity during early infection events was detected using an inverted confocal microscope (Nikon eclipse Ti in combination with a Yokogawa CSU-W1 spinning disk confocal system) and images were processed using ImageJ version 1.48 (US National Institutes of Health). GFP fluorescence was excited at 488 nm, while DsRed fluorescence was excited at 543 nm.

### Analysis of infection events, nodule number and nodule diameter

Composite plants expressing any of the different silencing, overexpressing, or control constructs were selected as described above. These roots were transferred to pots with vermiculite and inoculated with *R. tropici*–GUS to analyze IT progression, nodulation, and nitrogen fixation. Infection events were analyzed in the control and *PvCRT08* transgenic roots under a Zeiss Axioskop light microscope (Carl Zeiss, Jena, Germany) equipped with a X63 objective. Images were captured by a Nikon Coolpix 5000 camera with a UR-E6 adapter and an MDC lens attached to the microscope. GUS activity was analyzed according to the method of Jefferson **[25]**. Images of nodulated transgenic roots at 7, 10, 13, and 21 dpi stained with GUS were taken using a Perfection 4490 scanner (Epson) and captured in TIFF format at a resolution of 6108 × 6108 pixels. Nodule diameter was measured using ImageJ 1.48 (US National Institutes of Health) and classified according to their diameter (d) into four groups: Group I (d < 0.5 mm), Group II (0.5 < d ≤ 1.0 mm), Group III (1.0 < d ≤ 1.5 mm), and Group IV (1.5 < d <2.0 mm). The number of nodules was counted manually at 21 dpi.

### Acetylene reduction analysis

The acetylene reduction assay **[26]** was used to quantify the nitrogenase activity in transgenic nodules at 21 dpi. The nodulated roots were transferred to bottles with rubber seal stoppers and acetylene was injected into each bottle until a final concentration of 10% of the gas phase was reached. Each sample was incubated for 120 min at room temperature, and ethylene production was determined by gas chromatography in a Variant model 3300 chromatograph. Specific activity is expressed as µmol ethylene^−1^·(g nodule dry weight)^−1^·h^−1^

### Statistical analysis

Statistical analyses were computed using GraphPad Prism version 6.00 for Windows, (GraphPad Software, San Diego, CA, USA). Significance tests were performed using an unpaired Student’s t-test. Differences were considered significant if p < 0.05. Results are presented as means ± standard error of the mean.

## Results

### Calreticulin genes constitute a small family in the *P. vulgaris* genome

To examine the presumed participation of *P. vulgaris* calreticulin in its symbiotic interaction with rhizobia, we first identified the calreticulin genes in the *P. vulgaris* genome by searching the Phytozome v12 database (http://phytozome.jgi.doe.gov/pz/portal.html) **[19]**, using Phvul.007G200800, PvCRT08 (previously identified in a phosphoproteomic analysis, Ortega-Ortega, unpublished results), as query. Three calreticulin genes were identified in the genome of *P. vulgaris*, similar to data reported in monocotyledons and non-leguminous dicotyledons, and consistent with those found in the legume *G. max* (**S3 Table**). The deduced calreticulin proteins were named by including two digits of their access number: PvCRT08, PvCRT25 and PvCRT53 **[12,27,28]**. Analysis of the exon-intron organization showed that the gene structure is highly conserved among *PvCRT* genes, with 12 to 14 exons, which is a similar exon organization reported in *G. max* (**S1 Fig**) and *M. truncatula* **[29]**. The nucleotide sequence identity among *PvCRT* genes ranged from 59.7% to 69.8% (**S4 Table**), while identity between deduced amino acid sequences of PvCRT ranged from 50.1% to 80% (**S5 Table**). Multisequence alignment of protein sequences deduced from PvCRT and Arabidopsis AthCRT revealed the conserved residues and motifs previously reported by Michalak et al. **[13]** (**S2 Fig**). In addition, the PSORT program predicted that PvCRT proteins are located mainly i n peroxisome, lysosome, nucleus, and endoplasmic reticulum; and their theorical molecular weights range from 48 to 53 kDa (**S6 Table**).

To gain further insight into the evolutionary relationship between PvCRT proteins, a phylogenetic analysis was performed using sequences from both legumes (*G. max, M. truncatula, P. vulgaris)* and non-legumes (*O. sativa, A. thaliana*, *M. polymorpha, S. moellendorffii*) (**S1 Table**). In relation to the *A. thaliana* CRT family, the proteins clustered phylogenetically into two clades (**S3 Fig**), whereas PvCRT08 clustered into a subclade closely related to AthCRT1 and AthCRT2, which showed high expression in root tips, floral tissues and around leaf vascular tissues **[27]**.

### *PvCRT08* transcript is highly expressed in roots and after inoculation with rhizobia

*CRT* genes have been reported to show tissue-specific expression patterns in different plant systems and during various stages of development **[28,30]**. To determine *PvCR*T transcript levels, qPCR analysis was performed on different *P. vulgaris* tissues, such as root hairs, root apices, and stripped roots. As shown in **Fig 1**, the accumulation of *PvCRT* transcripts showed different patterns in the tissues examined. The expression pattern of *PvCRT53* could hardly be detected in any tissue, while *PvCRT25* showed the highest relative accumulation in stripped roots. However, the abundance of *PvCRT08* transcripts was the highest in all tissues tested.

**Fig 1.**
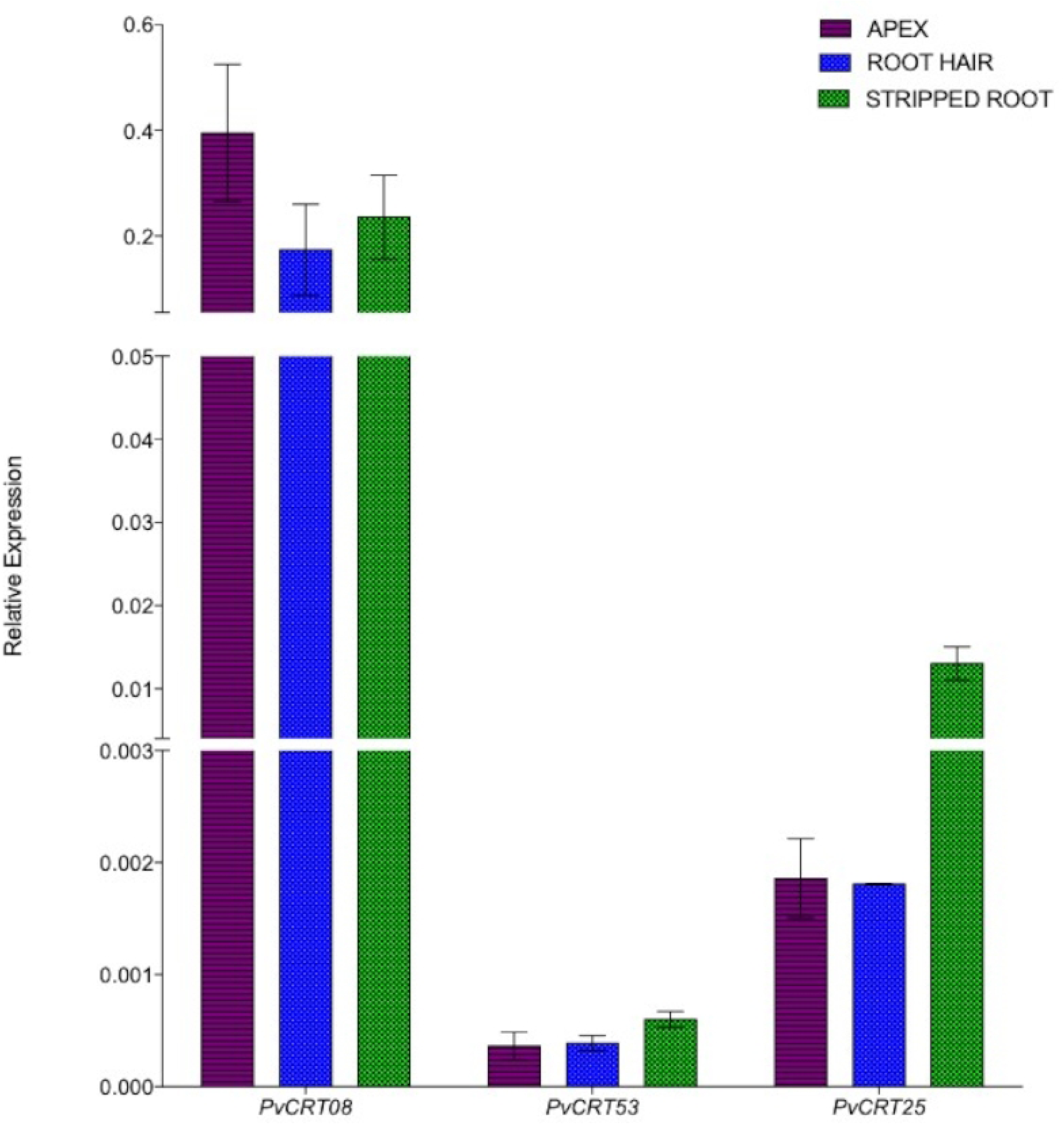
*PvCRT08* transcript is highly expressed in roots. Transcript accumulation profiles of *P. vulgaris* calreticulin genes in root hairs, root tips, and bare roots of seedlings harvested 2 days post-germination. Total RNA was isolated from each biological sample. The elongation factor (EF1α) was used as endogenous reference gene. Bars represent the mean ± SEM of three biological replicates (n > 6) with three technical replicates each.

These results are consistent with the *in silico* analysis based on RNA-sequencing, reported in the *P. vulgaris* Gene Expression Atlas (PvGEA, http://plantgrn.noble.org/PvGEA/). Interestingly, the *PvCRT53* gene was expressed to a greater extent in the pods and leaf tissues collected 5 and 21 days after plant inoculation with rhizobium. In contrast, no *PvCRT25* transcripts were detected in any of the organs or tissues analyzed (**S4A Fig**). Consistent with the PvGEA data, the *PVCRT08* gene was expressed in most of the organs and tissues examined and its transcript levels increased in inoculated roots and prefixed nodules, collected 5 dpi (**S4B Fig**). Furthermore, our previous findings indicate that the relative abundance of *PvCRT08* increased after Nod factor treatment (Ortega-Ortega, unpublished data). Taken together, these results strongly suggest that *PvCRT08* is involved in the symbiosis of the common bean root with *R. tropici;* therefore, we decided to characterize it functionally using reverse genetics.

### *PvCRT08* is expressed during different stages of nodulation with *R. tropici*

To investigate whether the abundance of *PvCRT08* transcripts varies after inoculation with *R. tropici* CIAT899, we monitored *PvCRT08* mRNA levels by RT-qPCR in whole roots during the early stages of nodulation (**Fig 2A**). In addition, the relative abundance of *PvCRT08* transcripts was analyzed in nodules and in nodule-free roots at 7, 10, 14 and 21 dpi (**Fig 2B**). During the early stages, *PvCRT08* transcript levels increased at 7 dpi, in whole roots, while during late stages *PvCRT08* transcript levels were significantly higher in nodules at 7, 10 and 21 dpi, and decreased at 14 dpi compared to nodule-free roots.

**Fig 2.**
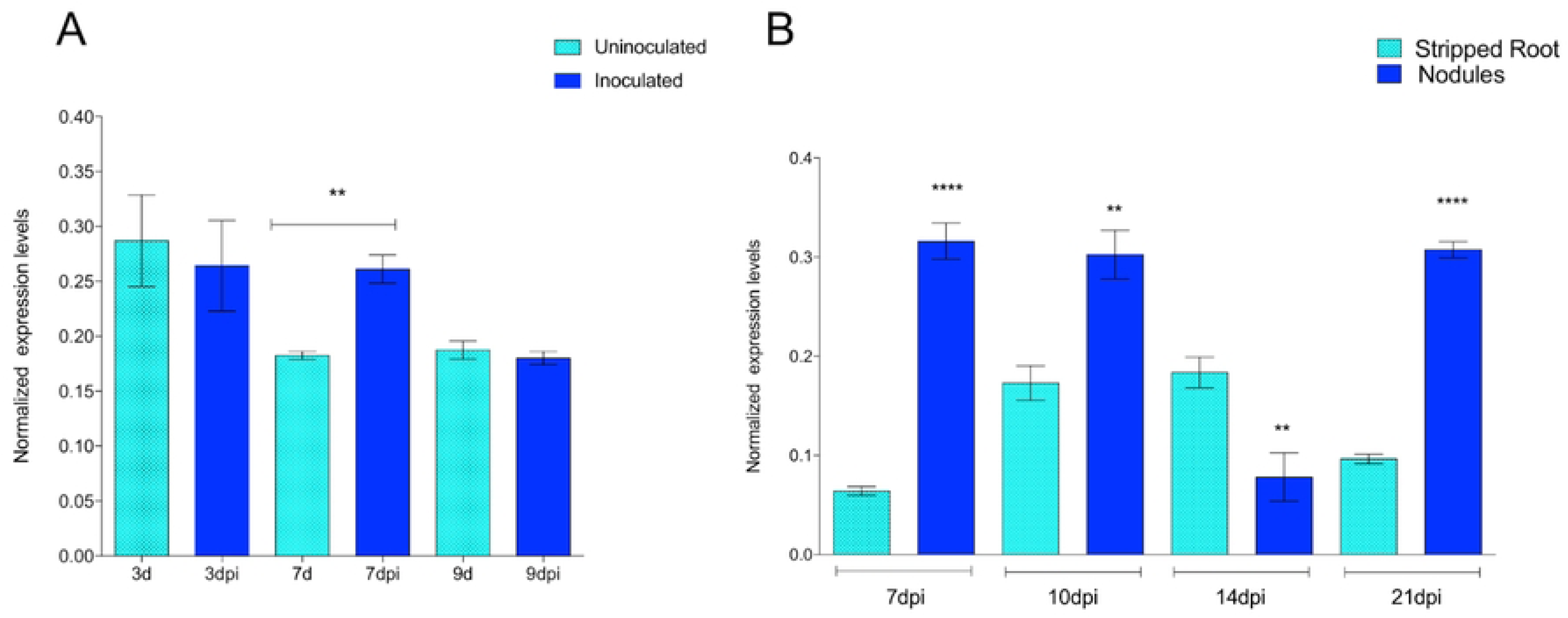
*PvCRT08* transcript levels increased during nodule development. (A) Accumulation of *PvCRT08* transcripts in whole roots that were either uninoculated or inoculated with *R. tropici* CIAT899, at the indicated number of days post-inoculation (dpi). (B) Relative accumulation of *PvCRT08* transcripts in nodules and nodule-free roots at the indicated number of days post-inoculation (dpi). For both A and B, total RNA was isolated from each biological sample and used in RT-qPCR analysis. The elongation factor EF1α was used as an endogenous reference gene. Each bar represents the mean ± SEM of two independent biological replicates (n> 6) with three technical repeats. \*\**p*<0.005 and \*\*\*\**p*<0.0001 based on Student’s *t*-test.

### Activity of the *PvCRT08* promoter was detected in root hairs, cortical cells, and in infected and vascular bundles of mature nodules

To examine the spatial and temporal gene expression of the *PvCRT08* gene, a transcriptional fusion of the *PvCRT08* promoter (2086 pb region upstream of the start codon) with the chimeric reporter GFP-GUS was constructed. Transgenic roots of *P. vulgaris* were generated by *R. rhizogenes* K599-mediated transformation with p*PvCRT08*:GFP-GUS. *PvCRT08* promoter activity was monitored by measuring GUS activity or GFP fluorescence in whole roots, hairy roots and nodules of the composite plants **[16]**. The spatial expression pattern of *PvCRT08* showed basal activity in vascular bundles, root tips, emerging primordial roots, and uninoculated root hairs (**Fig 3A to 3F**). Promoter activity was also monitored in transgenic roots after inoculation with *R.tropici* DsRed. In inoculated roots, promoter activity was detected in root hairs at 3 and 5 dpi, mainly in root hairs harboring growing ITs (**Fig 4**) and in adjacent dividing cortical cells (**Figure 3G**). During nodule organogenesis, high p*PvCRT8*:GFP-GUS activity was found in the core tissue of nodule-forming primordia, at 7 dpi (**Fig 3H**) and in infected cells and vascular bundles of mature nodules at 21 dpi (**Fig 3I and 3J**). This expression pattern suggests that *PvCRT08* has an important role during lateral root development, the progression of infection threads, as well as during nodule organogenesis.

**Fig 3.**
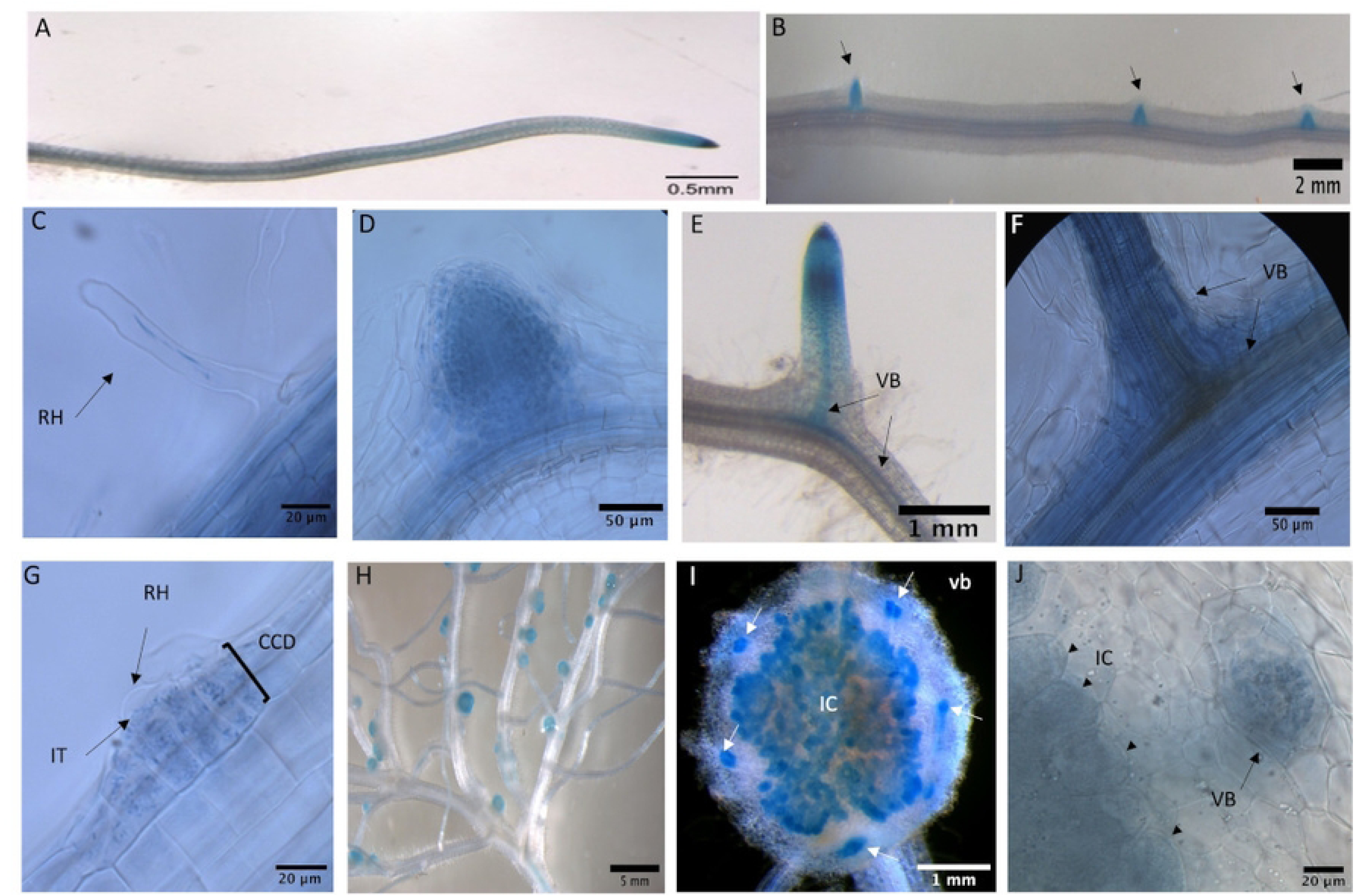
Promoter activity of *PvCRT08* was detected in uninoculated roots and during nodule organogenesis. Promoter activity was detected in the vascular bundles of uninoculated *P. vulgaris* roots (A), and in emerging lateral root primordia (B, D, E, F). Promoter activity was also detected in root hairs of uninoculated roots (C). The spatiotemporal expression pattern of the *PvCRT08* promoter was monitored in transgenic roots after-inoculation with *R. tropici* DsRed (G–J). (G) Infection site at 5 dpi. (H) Nodule primordium at 7 dpi. (I and J) Mature nodules at 21 dpi. (J) Free-hand sections of nodules at 21 dpi. RH: root hair; IT: infection thread; CCD: cortical cell division; IC: infected cell.

**Fig 4.**
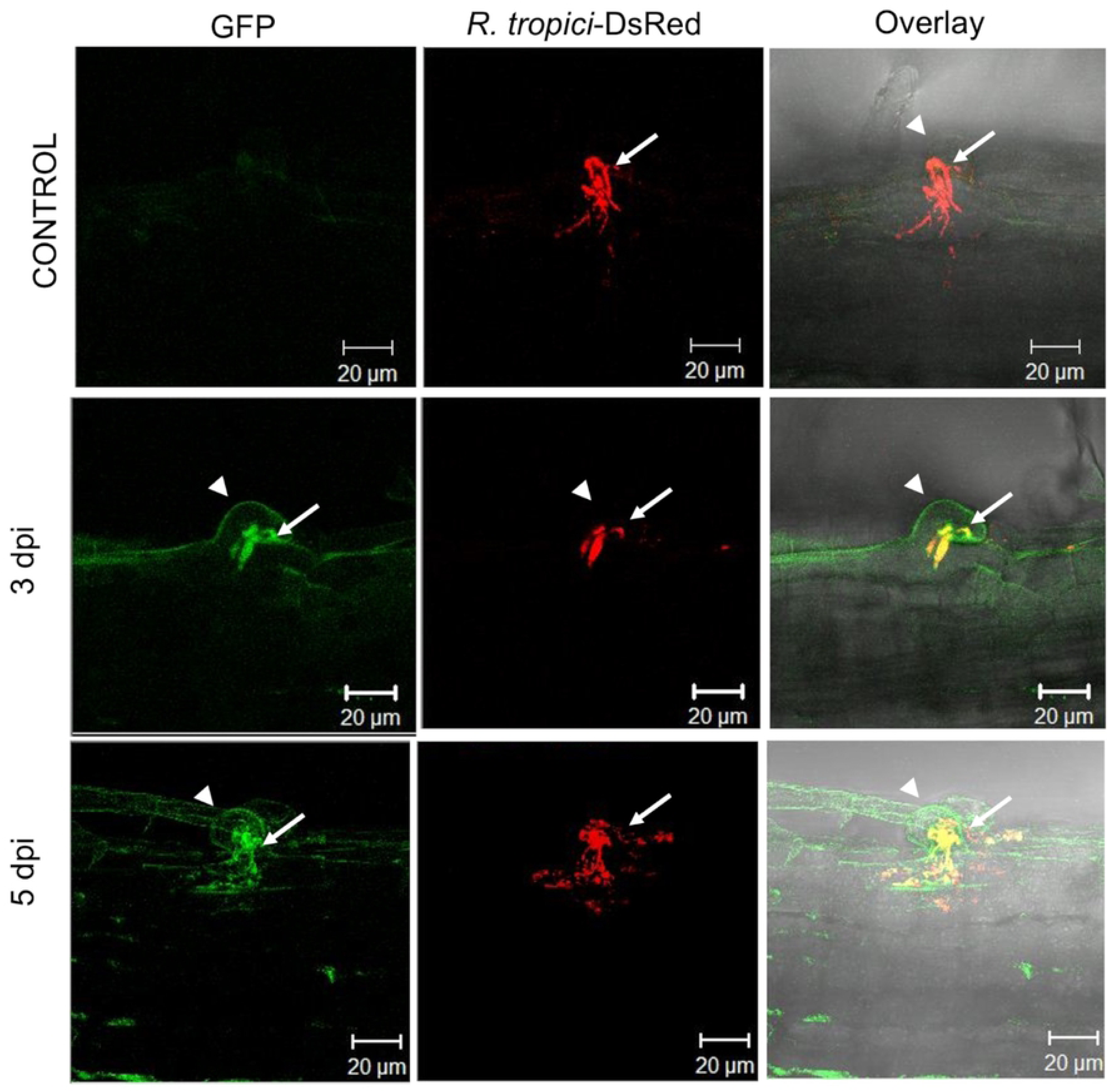
Activity of the *PvCRT08* promoter was identified in root hairs containing growing ITs. Promoter activity was monitored in root hairs after inoculation with *R. tropici*-DsRed at 3 and 5 dpi. Control transgenic roots harboring a vehicle without promoter sequence. Promoter activity was observed by confocal microscopy on transgenic roots expressing 2086 pb of the *PvCRT08* promoter region fused with GFP and GUS. dpi: days post-inoculation. Arrows indicate infection threads and the arrowhead points out a root hair.

### Loss-of function of the *PvCRT08* gene increases the number of infection events and nitrogen fixation levels

To assess the role of *PvCRT08* in nodulation, the effects of RNAi-mediated downregulation of this gene in *P. vulgaris* roots were investigated. To achieve this goal, transgenic bean roots were generated using *R. rhizogenes* K599 carrying *PvCRT08*-RNAi, and the control vector (roots transformed with a vector expressing an RNAi construct harboring an irrelevant sequence). Transgenic roots were identified by detecting red fluorescence derived from the red fluorescent protein marker (TdTomato) **[23]**. The abundance of *PvCRT08* transcripts in *PvCRT08* knockout roots was 76% lower compared to the control transgenic roots. This effect was gene-specific, as no statistically significant differences were detected in the abundance of *PvCRT2* and *PvCRT53* transcripts in the transgenic *PvCRT08*-RNAi roots compared to the control (**S5 Fig**).

To evaluate the participation of *PvCRT08* during the symbiotic relationship, the number of infection events within cortical cells of silenced roots inoculated with *R. tropici* CIAT899 was examined. As shown in **Fig 5**, the number of infections within cortical cells of silenced roots at 7 dpi was significantly higher than in control roots. These results suggest that *PvCRT08* affects the number of infections during the early stages of symbiosis.

**Fig 5.**
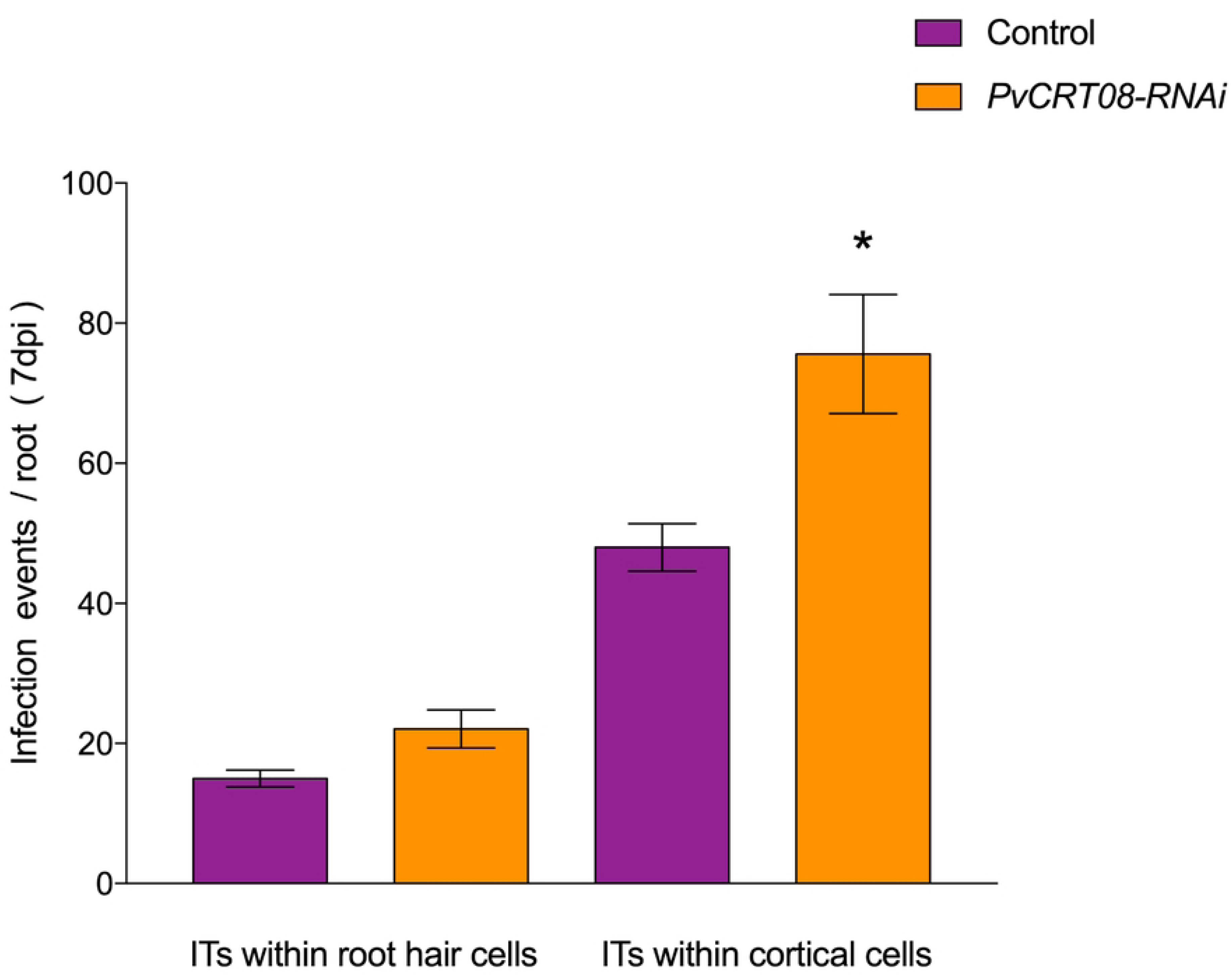
*PvCRT08*-RNAi increased the numbers of infection events in composite plants. Number or infection events within root hairs or cortical cells of *PvCRT08*-RNAi and control transgenic roots at 7 dpi with *R. tropici*. Values are mean + SEM with n>9 roots for each condition * p<0.05 according to Student’s t-test.

To better understand the regulatory role of *PvCRT08* in early rhizobial symbiosis, we quantified the abundance of *PvCyclin*, *PvEnod2* and *PvNIN* transcripts in transgenic roots inoculated with *R. tropici* CIAT899 (3 and 7 dpi), as markers of symbiosis progression **(S6A, S6B, S7A, S7B, 8SA and 8SB Fig).** Cyclin genes are suitable markers of dividing cells such as nodule formation **[31]**. We did not detect statistical differences in *PvCyclin* transcript abundance in *PvCRT08*-RNAi infected roots at any of the times points compared to control roots **(S6A and S6B Fig).** However, *PvEnod2* transcript abundance, a well-established marker of cortical cell division during early nodulation **[32]**, was significantly decreased in *PvCRT08*-RNAi roots at 3 dpi, while at 7 dpi they showed increased transcript abundance compared to control roots (**S7A and S7B Fig**). *NIN* is a marker for IT formation **[33]**. Interestingly, *PvNIN* transcription levels were significantly reduced in roots lacking *PvCRT08* at 3 dpi, whereas no differences in transcript abundance were observed at 7 dpi compared with control roots (**S8A and S8B Fig**). These results indicate an impaired activation of the nodulation signaling pathway.

Since downregulation of *PvCRT08* increased IT formation and progression, it could be predicted that the number of nodules would also be higher in silenced roots compared to control roots. However, the results obtained indicate that there were no significant differences in the total number of nodules in *PvCRT08*-RNAi roots (39.08 ± 2.32) compared to control transgenic roots (42.32 ± 2.94) at 21 dpi (**Fig 6A**). Interestingly, the nodules in the *PvCRT08*-RNAi roots were significantly larger than those in the control roots (**S9 Fig**) and were more efficient at nitrogen fixation (45%) compared to the control at 21 dpi (**Fig 6B**). These results reveal that *PvCRT08* plays an important role in nodule function.

**Fig 6.**
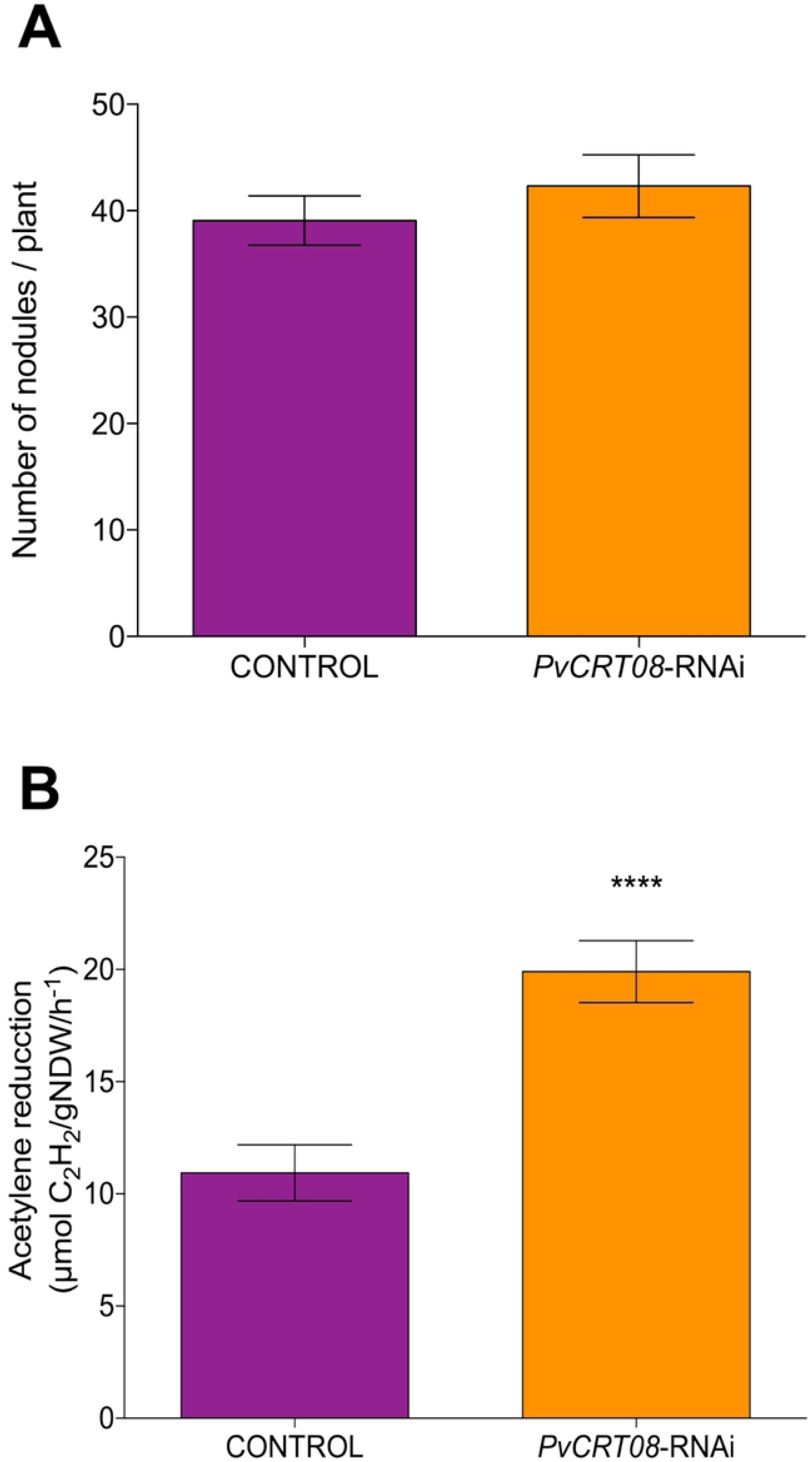
Silencing of *PvCRT08* does not affect the number of nodules but enhances nitrogen fixation capacity. Nodulation and nitrogen fixation capacity of *PvCRT08*-RNAi and control transgenic roots. **(A)** Nodules were collected and counted on *PvCRT08*-RNAi and control transgenic roots inoculated with *R. tropici* CIAT-899 at 21 dpi. **(B)** Nitrogenase activity was determined by acetylene reduction in nodules collected from *PvCRT08*-RNAi and control transgenic roots at 21 dpi. Bars represent the mean ± SEM for three biological replicates with *n*=10. \*\*\*\**p*<0.0001 based on Student’s *t-*test.

### Overexpression of *PvCRT08* disrupts infection progression, decreases nodule formation, and negatively affects nitrogen fixation efficiency

To analyze the effect of *PvCRT08* overexpression during nodulation, the transcript levels of *PvCRT08* in *P. vulgaris* composite plants were overexpressed, using the 35S promoter (**S10 Fig**). Subsequently, the levels of the gene transcript were determined in the overexpressing transgenic roots, revealing that the gene was expressed 3.5-fold higher (p < 0.001, *Student’s t*-test) than in control transgenic roots (containing an empty pH7WG2tdT vector (24). Following overexpression of *PvCRT08*, 52% of infection were arrested at the base of the root hairs at 7 dpi (**Fig 7**), whereas 87% of infections reached the cortical cells in the control transgenic roots. Unlike to the results for *PvCRT08*-RNAi, the abundance of *PvCyclin* and *PvNIN* transcripts in overexpressed transgenic roots decreased significantly at 3 and 7 dpi. In contrast, *PvCRT08* overexpression led to a marked upregulation of *PvEnod2* transcripts at 3 dpi (**S7C Fig**).

**Fig 7.**
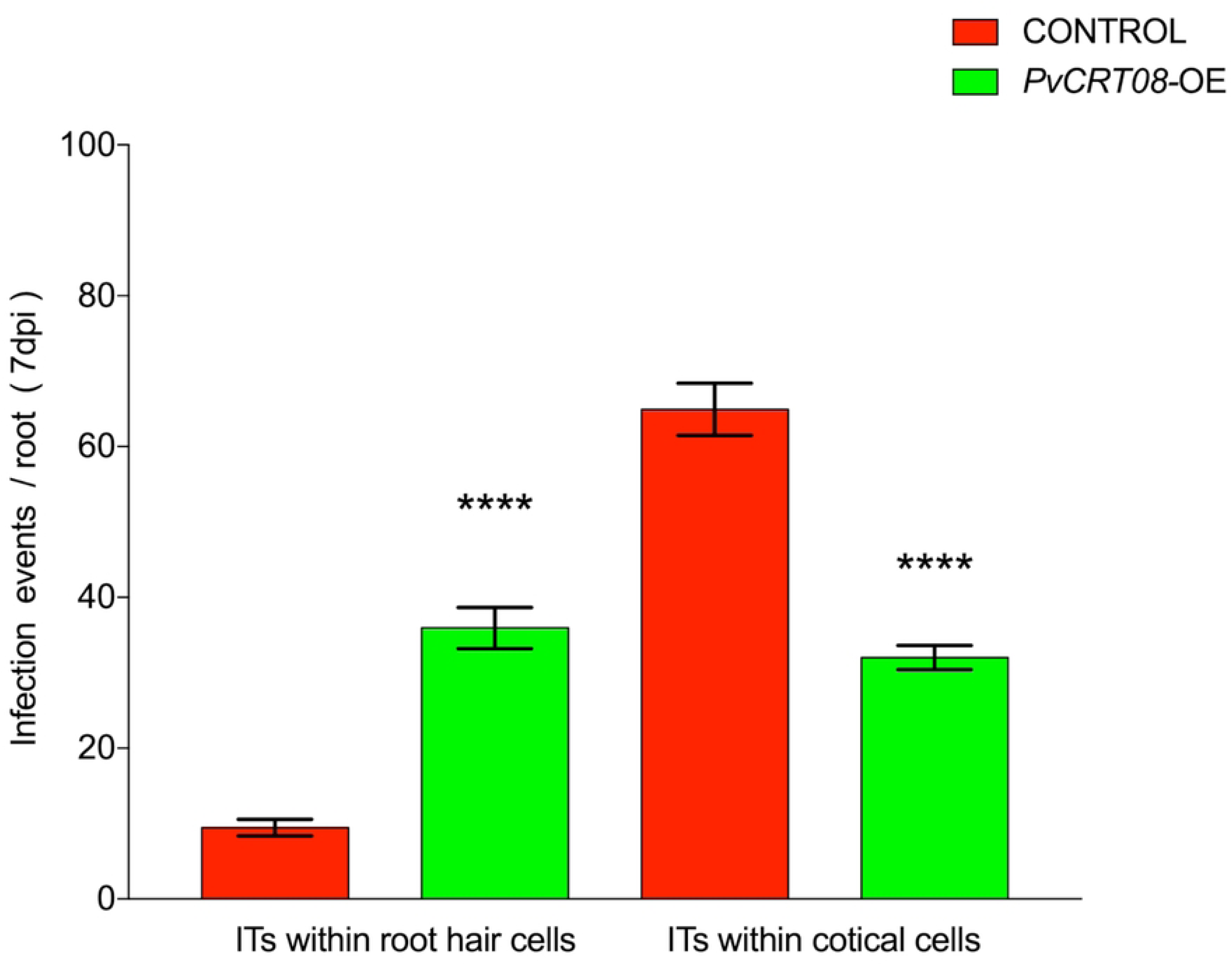
Overexpression of *PvCRT08* significantly reduces the number of infection events. Number of infection events within root hairs or cortical cells of transgenic *PvCRT08*-OE and control roots at 7 dpi with *R. tropici*. Values are expressed as mean + SEM with n>9 roots for each condition **** p<0.0001 according to Student’s t-test

In addition, the average number of nodules per plant in *PvCRT08*-OE was 54% lower than the control at 21 dpi, while they showed a nitrogen fixation capacity 21% lower than that of the control roots (**Fig 8**). Interestingly, no differences were observed in the total size of the nodules in the overexpressed roots (**S11 Fig**). These results indicate that PvCRT08 affects nodule development and function.

**Fig 8.**
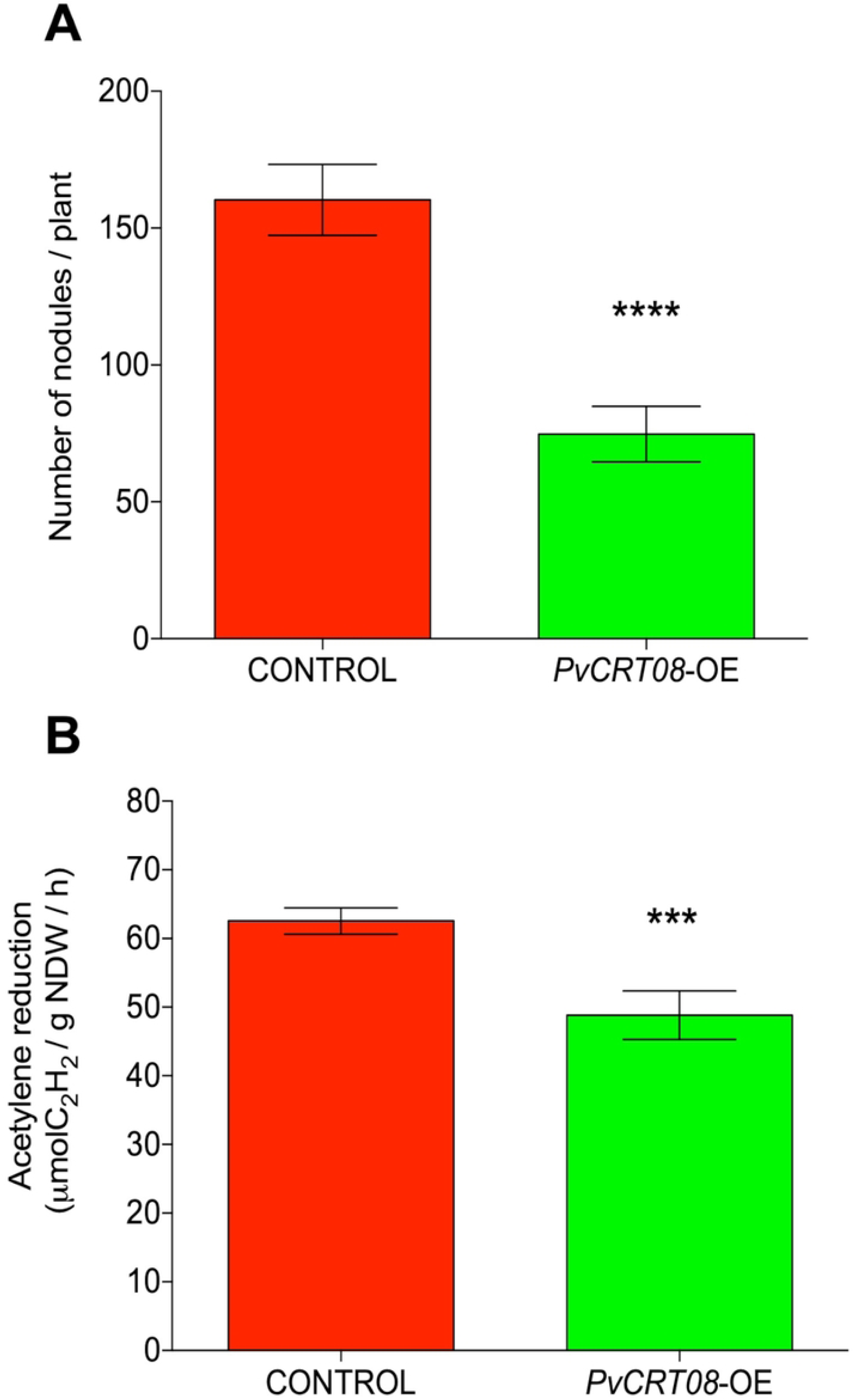
Overexpression of *PvCRT08* negatively affects the number of nodules and nitrogen fixation capacity. Nodulation and nitrogen-fixing capacity of *PvCRT08*-OE transgenic and control roots. **(A)** Nodules were collected and counted from control and *PvCRT08*-OE transgenic roots inoculated with *R. tropici* CIAT-899 at 21 dpi. **(B)** Nitrogenase activity determined by acetylene reduction in nodules collected from *PvCRT08*-OE and control transgenic roots at 21 dpi. Bars represent the mean ± SEM of four biological replicates with *n*=10.\*\*\*\**p*<0.0001, *** *p*<0.001, based on Student’s *t-*test.

## Discussion

Calreticulins are multifunctional proteins involved in calcium homeostasis, protein folding, and cellular signaling **[12,13]**. In legumes, similar functions of calreticulins have been described during plant–microbe interactions **[34]**. For instance, in *M. truncatula*, CRT is essential for calcium mobilization and the accommodation of *Rhizophagus* within cortical cells, as well as for arbuscule development **[35]**. The results presented here provide compelling evidence that *PvCRT08* plays a similar dual role in regulating early infection events and subsequent nodule development and function in *P. vulgaris*. In mycorrhizal roots of *M. truncatula*, colocalization of CRT and calcium ions were found on the surface of fungal structures and at the periphery of infected cortex cells **[35]**. The promoter activity of *PvCRT08* was detected in root hairs and infection sites after inoculation with *R. tropici,* suggesting that *PvCRT08* could increase Ca^2+^ ions flux.

The downregulation of *PvCRT08* altered the transcriptional responses of key nodulation genes such as *PvNIN* and *PvEnod2*, which are essential for infection thread progression and cortical cell division in the early phases of nodule development **[32,33]**. Furthermore, *PvCyclin* and *PvNIN* levels in overexpressed transgenic roots decreased significantly at 3 and 7 dpi. This suggests that *PvCRT08* may act as a molecular checkpoint to optimize symbiotic outcomes during early stages. This activity resembles the expression pattern of *NIN*, *ENOD11* and the activated form of Ca^2+^/calmodulin-dependent protein kinase (CCaMK), which are required not only for the formation of the infection thread, but also for maintaining the development and function of the nodule **[36,37]**.

Silencing *PvCRT08* increased the number of infection threads and enhanced nitrogen fixation efficiency, while its overexpression impaired infection progression and reduced both nodulation and symbiotic effectiveness. The contrasting phenotypes observed in *PvCRT08*-silenced lines versus overexpressed lines highlight the importance of maintaining optimal CRT activity. Similar regulatory patterns have been observed in CCaMK, where specific deletion of the autoinhibition domain -that negatively regulates the kinase activity- leads to the autoactivation of the nodulation signaling pathway in the *Sinorhizobium meliloti–M. truncatula* symbiosis **[36]**. Notably, overexpression of the His173 mutant of CRT2 in Arabidopsis confers enhanced resistance to PstDC3000 infection **[38]**. Because this conserved histidine is critical for CRT chaperone function **[39]**, it is plausible to propose that *PvCRT08* influences infection progression through its chaperone activity.

It is noteworthy that the increased nitrogenase activity in larger nodules of *PvCRT08*-silenced roots, suggests that this gene protein product also influences the metabolic and physiological competence of the nodules, extending its function beyond early signaling events. CRT, predominantly located in the endoplasmic reticulum, has been shown to localize in plasmodesmata connecting uninfected cells in nitrogen-fixing root nodules of the Australian tree *Casuarina glauca* **[34]**. Plasmodesmal connections between infected cells, and to a lesser extent between infected and uninfected cells, were reduced during the differentiation of infected cells, suggesting a their role in cell-to-cell transport **[40]**. We propose that, during nodule development in *P. vulgaris*, infected cells reduce their plasmodesmatic connections with neighboring cells, a process linked to the decrease of *PvCRT08* and which probably facilitates a greater exchange of metabolites.

Taken together, our findings highlight *PvCRT08* as a central regulator of symbiotic efficiency in the common bean, operating at multiple stages of the infection and nodulation processes. The dual impact of its downregulation and overexpression underscores the need for a tight controlled balance of CRT activity, which may serve as a molecular strategy to optimize the equilibrium between infection efficiency and nodule functionality.

## Supplementary Figure captions

**S1 Fig**. Exon-intron organization of *CRT* genes

**S2 Fig.** Alignment of the deduced amino acid sequences of PvCRTs and AthCRTs.

**S3 Fig**. Phylogenetic tree of CRT family proteins

**S4 Fig.** Transcript abundance of *CRT* genes in different organs and tissues of *P. vulgaris*.

**S5 Fig.** Expression of *PvCRT* genes in control and *PvCRT08*-RNAi transgenic roots at 10 days post emergence

**S6 Fig**. Transcript abundance of *PvCyclin* in CTR-RNAi transgenic roots at 3 dpi (A) and 7 dpi (B). Transcript abundance of *PvCyclin* in CTR overexpression roots a 3 dpi (C) and 7 dpi (D).

**S7 Fig**. Transcript abundance of *PvEnod2* in CTR-RNAi transgenic roots at 3 dpi (A) and 7 dpi (B). Transcript abundance of *PvEnod2* in CTR overexpression roots a 3 dpi (C) and 7 dpi (D).

**S8 Fig**. Transcript abundance of *PvNIN* in CTR-RNAi transgenic roots at 3 dpi (A) and 7 dpi (B). Transcript abundance of *PvNIN* in CTR overexpression roots a 3 dpi (C) and 7 dpi (D).

**S9 Fig**. Nodule diameters on *PvCRT08*-RNAi and control transgenic roots after inoculation with *R. tropici* expressing GUS

**S10 Fig**. Expression of *PvCRT* genes in control and *PvCRT08*-OE transgenic roots at 10 days post emergence.

**S11 Fig**. Nodule diameters on *PvCRT08*-OE and control transgenic roots after inoculation with *R. tropici* expressing GUS

## Supplementary Table captions

**S1 Table.** Calreticulin protein annotations from multiple plant genomes and human

**S2 Table**. Gene-specific oligonucleotides

**S3 Table.** Size of the *CRT* gene family in various plants

**S4 Table.** Percentage of nucleotide sequence identity among *P. vulgaris CRT* genes

**S5 Table**. Percentage of amino acid sequence identity between PvCRT and AthCRT proteins

**S6 Table.** Characteristics of predicted bean calreticulin proteins.

## Notes

### Competing Interest Statement

The authors have declared no competing interest.

## References

1. Abd-Alla MH, Al-Amri SM, El-Enany AWE. Enhancing Rhizobium–Legume Symbiosis and Reducing Nitrogen Fertilizer Use Are Potential Options for Mitigating Climate Change. Agriculture. 2023; 13: 2092.

2. Goyal RK, Mattoo AK, Schmidt MA. Rhizobial–Host Interactions and Symbiotic Nitrogen Fixation in Legume Crops Toward Agriculture Sustainability. Frontiers in Microbiology. 2021; 12.

3. Mus F, Crook MB, Garcia K, Garcia Costas A, Geddes BA, Kouri ED, et al. Symbiotic Nitrogen Fixation and the Challenges to Its Extension to Nonlegumes. Applied and environmental microbiology. 2016; 82(13): 3698–3710.

4. Smercina DN, Evans SE, Friesen ML, Tiemann LK. To Fix or Not To Fix: Controls on Free-Living Nitrogen Fixation in the Rhizosphere. Applied and environmental microbiology. 2019; 85(6): e02546–18.

5. Gepts P. Phaseolus vulgaris (beans). In Brenner S, Miller JH. Encyclopedia of genetics.: Academic Press; 2001. p. 1444–1445.

6. Shumilina J, Soboleva A, Abakumov E, Shtark OY, Zhukov VA, Frolov A. Signaling in Legume–Rhizobia Symbiosis. International Journal of Molecular Sciences. 2023; 24(24): 17397.

7. Yang J, Lan L, Jin Y, Yu N, Wang D, Wang E. Mechanisms underlying legume–rhizobium symbioses. Journal of Integrative Plant Biology. 2022; 64(2): 244–267.

8. de Carvalho-Niebel F, Fournier J, Becker A, Arancibia MM. Cellular insights into legume root infection by rhizobia. Current opinion in plant biology. 2024; 81: 102597.

9. Porter SS, Dupin SE, Denison RF, Kiers ET, Sachs JL. Host-imposed control mechanisms in legume–rhizobia symbiosis. Nature Microbiology. 2024; 9(8): 1929–1939.

10. Zartdinova R, Nikitin A. Calcium in the life cycle of legume root nodules. Indian Journal of Microbiology. 2023; 63(4): 410–420.

11. Zhang X, Wu J, Kong Z. Cellular basis of legume–rhizobium symbiosis. Plant Communications. 2024; 5(11).

12. Jia XY, He LH, Jing RL, Li RZ. Calreticulin: conserved protein and diverse functions in plants. Physiologia plantarum. 2009; 136(2): 127–138.

13. Michalak M, Groenendyk J, Szabo E, Gold LI, Opas M. Calreticulin, a multi-process calcium-buffering chaperone of the endoplasmic reticulum. The Biochemical Journal. 2009; 417(3): 651–666.

14. Thelin L, Mutwil M, Sommarin M, Persson S. Diverging functions among calreticulin isoforms in higher plants. Plant signaling & behavior. 2011; 6(6): 905–910.

15. Navazio L, Miuzzo M, Royle L, Baldan B, Varotto S, Merry AH, et al. Monitoring endoplasmic reticulum-to-Golgi traffic of a plant calreticulin by protein glycosylation analysis. Biochemistry. 2002; 41(48): 14141–14149.

16. Estrada-Navarrete G, Alvarado-Affantranger X, Olivares JE, Guillen G, Diaz-Camino C, Campos F, et al. Fast, efficient and reproducible genetic transformation of Phaseolus spp. by Agrobacterium rhizogenes. Nat Protoc. 2007; 2(7): 1819–1824.

17. Young JM, Kuykendall LD, Martínez-Romero E, Kerr A, Sawada H. A revision of Rhizobium Frank 1889, with an emended description of the genus, and the inclusion of all species of Agrobacterium Conn 1942 and Allorhizobium undicola de Lajudie et al. 1998 as new combinations: Rhizobium radiobacter, R. rhizogenes, R. rubi. International journal of systematic and evolutionary microbiology. 2001; 51(Pt 1): 89–103.

18. Broughton WJ, Dilworth MJ. Control of leghaemoglobin synthesis in snake beans. Biochem J. 1971; 125(4): 1075–1080.

19. Goodstein DM, Shu S, Howson R, Neupane R, Hayes RD, Fazo J, et al. Phytozome: a comparative platform for green plant genomics. Nucleic acids research. 2012; 40(D1): D1178–D1186.

20. Livak KJ, Schmittgen TD. Analysis of Relative Gene Expression Data Using Real-Time Quantitative PCR and the 2−ΔΔCT Method. Methods. 2001; 25(4): 402–408.

21. Schmittgen TD, Livak KJ. Analyzing real-time PCR data by the comparative CT method. Nat Protoc. 2008; 3(6): 1101–1108.

22. Karimi M, Inze D, Depicker A. GATEWAY vectors for Agrobacterium-mediated plant transformation. Trends Plant Sci. 2002; 7(5): 193–195.

23. Valdes-Lopez O, Arenas-Huertero C, Ramirez M, Girard L, Sanchez F, Vance CP, et al. Essential role of MYB transcription factor: PvPHR1 and microRNA: PvmiR399 in phosphorus-deficiency signalling in common bean roots. Plant Cell Environ. 2008; 31(12): 1834–1843.

24. Ortega-Ortega Y, Carrasco-Castilla J, Juárez-Verdayes MA, Toscano-Morales R, Fonseca-García C, Nava N, et al. Actin depolymerizing factor modulates rhizobial infection and nodule organogenesis in common bean. Int J Mol Sci. 2020; 21(6): 1970.

25. Jefferson RA. Assaying chimeric genes in plants: The GUS gene fusion system. Plant Molecular Biology Reporter. 1987; 5(4): 387–405.

26. Ramírez M, Valderrama B, Arredondo-Peter R, Soberón M, Mora J, Hernández G. Rhizobium etli Genetically Engineered for the Heterologous Expression of Vitreoscilla sp. Hemoglobin: Effects on Free-Living and Symbiosis. Molecular Plant-Microbe Interactions. 1999; 12(11): 1008–1015.

27. Christensen A,SK:TL, Zhang W, Tintor N, Prins D, Funke N, Michalak M, et al. Higher plant calreticulins have acquired specialized functions in Arabidopsis. PLoS One. 2010; 5(6): e11342.

28. Persson S, Rosenquist M, Svensson K, Galvão R, Boss WF, Sommarin M. Phylogenetic analyses and expression studies reveal two distinct groups of calreticulin isoforms in higher plants. Plant Physiology. 2003; 133(3): 1385–1396.

29. Wasąg P, Grajkowski T, Suwińska A, Lenartowska M, Lenartowski R. Phylogenetic analysis of plant calreticulin homologs. Molecular Phylogenetics and Evolution. 2019; 134: 99–110.

30. Denecke J, Carlsson LE, Vidal S, Höglund AS, Ek B, van Zeijl MJ, et al. The tobacco homolog of mammalian calreticulin is present in protein complexes in vivo. Plant Cell. 1995; 7(4): 391–406.

31. Jelenska J, Deckert J, Kondorosi E, Legocki, A.B.. Mitotic B-type cyclins are di erentially regulated by phytohormones and during yellow lupine nodule development. Plant Sci. 2000; 150: 29–39.

32. Stougaard J. Regulators and regulation of legume root nodule development. Plant Physiol. 2000; 124: 531–540.

33. Schauser L, Roussis A, Stiller J, Stougaard J. A plant regulator controlling development of symbiotic root nodules. Nature. 1999; 402(6758): 191–195.

34. Demchenko KN, Voitsekhovskaja OV, Pawlowski K. Plasmodesmata without callose and calreticulin in higher plants - open channels for fast symplastic transport? Front Plant Sci. 2014; 5: 74.

35. Sujkowska-Rybkowska M, Znojek E. Localization of calreticulin and calcium ions in mycorrhizal roots of Medicago truncatula in response to aluminum stress. Journal of Plant Physiology. 2018; 229: 22–31.

36. Gleason C, Chaudhuri S, Yang T, Muñoz A, Poovaiah BW, Oldroyd GED. Nodulation independent of rhizobia induced by a calcium-activated kinase lacking autoinhibition.. Nature. 2006; 441(7097): 1149–1152.

37. Marsh JF, Rakocevic A, Mitra RM, Brocard L, Sun J, Eschstruth A, et al. Medicago truncatula NIN is essential for rhizobial-independent nodule organogenesis induced by autoactive calcium/calmodulin-dependent protein kinase. Plant Physiology. 2007; 144(1): 324–335.

38. Qiu Y, Xi J, Du L, Roje S, Poovaiah BW. A dual regulatory role of Arabidopsis calreticulin-2 in plant innate immunity. The Plant Journal. 2012; 69(3): 489–500.

39. Guo L, Groenendyk J, Papp S, Dabrowska M, Knoblach B, Kay C, et al. Identification of an N-domain histidine essential for chaperone function in calreticulin. J Biol Chem. 2003; 278: 50645–50653.

40. Schubert M, Koteyeva NK, Zdyb A, Santos P, Voitsekhovskaja OV, Demchenko KN, et al. Lignification of cell walls of infected cells in Casuarina glauca nodules that depend on symplastic sugar supply is accompanied by reduction of plasmodesmata number and narrowing of plasmodesmata. Physiol Plant. 2013; 147: 524–540.

